# Family History of Depression is Associated with Alterations in Task-Dependent Connectivity between the Cerebellum and Ventromedial Prefrontal Cortex

**DOI:** 10.1101/851477

**Authors:** Lindsey J. Tepfer, Lauren B. Alloy, David V. Smith

## Abstract

**Background:** A family history of major depressive disorder (MDD) increases the likelihood of a future depressive episode, which itself poses a significant risk for disruptions in reward processing and social cognition. However, it is unclear whether a family history of MDD is associated with alterations in the neural circuitry underlying reward processing and social cognition.

**Methods:** We subdivided 279 participants from the Human Connectome Project into three groups: 71 with a lifetime history of MDD, 103 with a family history of MDD (FH), and 105 healthy controls (HC). We then evaluated task-based fMRI data on a social cognition and a reward processing task and found a region of the ventromedial prefrontal cortex (vmPFC) that responded to both tasks, independent of group. To investigate whether the vmPFC shows alterations in functional connectivity between groups, we conducted psychophysiological interaction (PPI) analyses using the vmPFC as a seed region.

**Results:** We found that FH (relative to HC) was associated with increased sadness scores, and MDD (relative to both FH and HC) was associated with increased sadness and MDD symptoms. Additionally, the FH group had increased vmPFC functional connectivity within the nucleus accumbens, left dorsolateral PFC, and subregions of the cerebellum relative to HC during the social cognition task.

**Conclusions:** These findings suggest that aberrant neural mechanisms among those with a familial risk of MDD may underlie vulnerability to altered social cognition.

## Introduction

Major Depressive Disorder (MDD) substantially interferes with the ability to carry out a normal life. As a leading contributor to global disability (Ferrari et al., 2013; World Health Organization, 2017), MDD carries profoundly disruptive symptoms that range from hopelessness (Abramson et al., 1989; Iacoviello et al., 2010) to diminished motivation (Sherdell et al., 2012; Treadway et al., 2012) as a part of anhedonia (American Psychiatric Association, 2013; Pizzagalli, 2014), and is a prominent predictor of suicidality (Klonsky et al., 2016; Nock et al., 2009). Symptom recurrence is likely (Kovacs et al., 2016; Solomon et al., 2000), and each subsequent episode increases the chance that more will follow (Colman et al., 2011; Hoertel et al., 2017). Depression is highly comorbid with substance use disorders (Swensden & Merikangas, 2000), including alcohol (Fergusson et al., 2009; Pedrelli et al., 2016; L. E. Sullivan et al., 2005) and tobacco (Covey et al., 1998; Weinberger et al., 2017), and there is evidence that the relationship between MDD and substance use may impart similar patterns in the children of those who suffer from depression (Luthar, Merikangas, Rounsaville et al., 1993) as both are highly familial (Merikangas &Swendsen, 1997). Importantly, the probability that an individual will experience MDD during their lifetime increases for those with a family history (FH) of depression (Klein et al., 2005, 2013; Monroe et al., 2014; Zimmermann et al., 2008). Indeed, those with depressed first-degree relatives are nearly three times as likely to develop the disorder later in life (Sullivan et al., 2000). Upon the onset of MDD, the systems that support social cognition also are affected (Cusi et al., 2012; Derntl et al., 2011), which limits the capacity to maintain supportive connections with others and negatively impacts recovery (George et al., 1989; Santini et al., 2015). Understanding the abnormalities among the neural mechanisms underlying social cognition and reward sensitivity in MDD, and how they are vulnerable in those with a familial risk of MDD, stands to elucidate the relationship between early pathophysiology in the brain and subsequently altered cognition.

A wealth of research suggests that many of the symptoms associated with MDD involve abnormalities in specific neural circuits. For example, studies have found grey matter volume reductions in frontotemporal areas (Cai et al., 2014; Serra-Blasco et al., 2013), the hippocampus (Arnone et al., 2013; Zhao et al., 2014), the orbitofrontal cortex (OFC; (Bremner et al., 2002; Lacerda et al., 2004), the ventromedial prefrontal cortex (vmPFC; (Wise et al., 2017), the dorsomedial prefrontal cortex (dmPFC), the anterior cingulate cortex (ACC), and the bilateral insula (Caetano et al., 2006; Salvadore et al., 2011). Recent reports also indicate a putative role for cerebellar grey matter volume decrease in depression (Wise et al., 2017). Indeed, abnormal structural circuitry in the cerebellum appears to underlie many common mental disorders (Romer et al., 2018) with a handful of studies indicating subregional specificity, such as the vermis and posterior cerebellum. For example, lesioning in the posterior and vermis subregions often is associated with emotion dysregulation and other affective symptoms (Schmahmann, 2004; Schmahmann et al., 2007), and increased anterior vermis volume has been found in MDD patients (Yucel et al., 2013). Likewise, white matter fractional anisotropy investigations also show disruptions across the lifespan among individuals with depression (Alexopoulos et al., 2008; Bae et al., 2006; Cruwys et al., 2014; Cullen et al., 2010; Steingard et al., 2002; Taylor et al., 2004; Yang et al., 2007). In conjunction with structural changes, MDD is associated with functional aberrations in systems associated with reward processing. Evidence of differences in activation and connectivity in healthy individuals relative to those with MDD typically implicate the OFC (Cheng et al., 2018; Ng et al., 2019), the ventral striatum (VS; (Furman et al., 2011; Kumar et al., 2018; Robinson et al., 2012), and the vmPFC (Koenigs et al., 2008; Myers-Schulz & Koenigs, 2012). Similarly, alterations in function also have been found in the cerebellum among depressed patients, including increased blood flow to the cerebellar vermis (Bench et al., 1992; Dolan et al., 1992), increasing vermis-PCC functional connectivity with symptom severity (Alalade et al., 2011), left-posterior cerebellar activation during rewarded sustained attention (Chantiluke et al., 2012), as well as decreases in vmPFC-bilateral lobule VIIb functional connectivity (Liu et al., 2012). Together, this work suggests that we may be able to understand some of the canonical symptoms seen in MDD symptomatology by taking a closer look at the neural correlates that underlie them.

Tied to the functional and structural impairments seen among reward processing systems (e.g., the striatum), a hallmark symptom of MDD involves the marked loss in the ability to experience pleasure (Cooper et al., 2018). Individuals with MDD often endure significant decreases in motivation (Hershenberg et al., 2016), diminishment in emotional responses to things that are pleasant (Alloy et al., 2016), and no longer pursue the interests they once found enjoyable (Fried & Nesse, 2014). Moreover, research implicates that social rewards share the neural circuitry represented by non-social rewards (Bhanji & Delgado, 2014; Fareri & Delgado, 2014). Thus, abnormalities in reward processing often overlap with social cognitive deficits, as reward system activation is critical for facilitating positive social exchanges (Carta et al., 2019; Smith & Delgado, 2015; Wang et al., 2016). Deficits in social cognition persist throughout recovery from MDD and are associated with later relapse (Inoue et al., 2004, 2006).

Individuals with a FH of depression also appear to be at risk for similar degrees of deviation in reward and social processing early on (Monk et al., 2008; Weinberg et al., 2015). For example, individuals with a familial risk of depression show abnormal responses to reward and punishment (McCabe et al., 2012), have reduced activity in the VS and ACC in response to social reward during adolescence (Olino et al., 2015), and decreased VS and anterior insula activity in response to non-social rewards during childhood (Luking et al., 2016). Furthermore, those with a familial history of MDD begin to show alterations in theory of mind (ToM) processing - the ability to infer the mental states of others (Harkness et al., 2011).

However, despite the work underscoring the relationship between reward, social cognition, and depression, it remains unclear whether individuals with a familial history of MDD demonstrate neural alterations associated with processing social stimuli in the absence of reward. Investigating how the neural and behavioral alterations observed in MDD emerge in association with a FH is key to understanding the mechanisms that drive these changes. Moreover, evaluating cases with a predisposition to such alterations may inform current treatment practices that can help provide early intervention upon the identification of at-risk populations. Thus, in the present study, we investigate whether a FH of depression, in the absence of a personal diagnosis, is associated with altered social cognition and reward processing in the brain. Our analyses focused on three main hypotheses. First, we expected to see increased uncertainty during ToM processing, as well as increased depressive symptoms, reports of sadness, alcohol, and tobacco use among individuals with a personal history of MDD, relative to those with a FH of MDD or HCs. Second, we predicted that the MDD group would show blunted temporoparietal junction (TPJ) response and increased vermis and posterior cerebellar activity during a ToM task, and decreased striatal activation in response to reward during a gambling task. Our third hypothesis was that the MDD group would show decreased connectivity between the cerebellum’s bilateral lobule VIIb and the vmPFC during the social task. For all of these hypotheses, we anticipated a dose-dependent type of effect across the three groups, in which the MDD group would show the highest magnitude of difference relative to the FH and HC groups, with the FH group resembling any effect seen in the MDD group, albeit at a smaller magnitude. For example, should the MDD group demonstrate hypothesized blunted TPJ activity during the ToM task, the FH group would similarly show decreases in TPJ activity relative to HCs, but not decreased to the same extent as the MDD group.

## Materials and Methods

### Participants

The sample included 279 participants (males, 120; females, 159; ages, 22-36; mean ± SD, 28.45 ± 3.75 years) selected from the Human Connectome Project (HCP) dataset, a large collection of primarily healthy young adults (N=1206) from the 1200 Subjects Release (WU-Minn HCP consortium S1200 Data). The study protocol was performed in compliance with the Code of Ethics of the World Medical Association (Declaration of Helsinki). Participants provided written informed consent, and procedures were approved by the ethics committee in accordance with guidelines of WU-Minn HCP. We subdivided participants into three groups. Individuals in the MDD group (N=71) were selected if they had a lifetime history of a DSM-IV Major Depressive Episode diagnosis. Individuals in the FH group (N=103) were selected if they had not received a DSM-IV Major Depressive Episode diagnosis in their lifetime, but either their mother or father had. We note that the 71 MDD and 103 FH comprised all participants who met criteria for these groups from the full 1206 HCP sample. HCs (N=105) consisted of participants who had not received a DSM-IV Major Depressive Episode diagnosis in their lifetime, and had no FH of psychiatric or neurologic disorders. The number of HCs that qualified based on these criteria was large (N=573); to facilitate comparison, we randomly selected HC participants to match our largest experimental group (N=105).

### Exclusion Criteria

Participants were excluded for head motion if their average relative head motion (calculated using a root-mean-squared approach; Jenkinson et al., 2002) exceeded the 75th percentile plus the value of 150% of the interquartile range of average relative head motion for all subjects (i.e., a standard boxplot threshold; (Smith et al., 2014). Additionally, participants who met DSM-IV criteria for alcohol abuse were excluded from the current study.

### Task Paradigms

All 279 participants in the current study performed 7 tasks in the scanner across two sessions, with two runs of each task. Here, we analyze data from only 2 tasks: social cognition and reward processing. Each task involved alternating blocks of stimuli. The descriptions of the two tasks of interest here are adaptations of the details provided elsewhere (Barch et al., 2013). The social cognition task required participants to indicate whether or not they interpreted a social interaction among brief animations of geometric shapes, a robust and reliable measurement of theory of mind processing. The reward processing task prompted participants to guess whether a ‘mystery card’ held a value of higher or lower than 5, and subsequently received feedback on the outcome of their choice. For more details regarding the block timing and composition of each task, see **Supplemental Information**.

### Image Acquisition

All 279 participants in the current study underwent T1 and T2-weighted structural scans, resting-state and task-based fMRI scans, and diffusion-weighted MRI scans, all via a 3T Siemens scanner with a standard Siemens 32 channel RF head coil. Identical multi-band EPI sequence parameters leveraged high spatial resolution at 2mm isotropic using the following parameters: repetition time (TR) = 720 ms; echo time (TE) = 33.1 ms, 72 slices, 2mm isotropic voxels, using a multi-band acceleration factor of 8 (Feinberg et al., 2010; Moeller et al., 2010; Setsompop et al., 2012; Xu et al., 2013). The current investigation used minimally preprocessed BOLD fMRI data to increase ease of use and mitigate the possibility of complex issues relating to imaging data quality; the general preprocessing steps are presented in full detail in previous reports (Glasser et al., 2013; Smith et al., 2013; Ugurbil et al., 2013; Van Essen et al., 2013). Tasks were performed twice, with one run in a left-to-right direction, and then reversed in the second run. In addition to the minimal preprocessing already featured in the HCP dataset, we identified and removed artifacts sourced from excess head-motion using the ICA-AROMA Software Package. Preceding ICA-AROMA, we first smoothed the data with a spatial smoothing of 6mm and the entire 4D dataset received grand-mean intensity normalization using a single multiplicative factor. This grand-mean intensity normalization step was applied to ensure scaling is not shifted following denoising and the normalization (mean = 10,000) is maintained. For convenience, the current study worked solely in volumetric MNI-space, although future work would benefit from transitioning to a CIFTI space (Coalson et al., 2018; Glasser, Coalson, et al., 2016; Glasser, Smith, et al., 2016>).

### fMRI Analysis

We analyzed task-based fMRI data using FEAT (FMRI Expert Analysis Tool) Version 6.00, part of FSL (FMRIB’s Software Library; (Jenkinson et al., 2012)). Specifically, we used FILM (FMRIB’s Improved Linear Model) with local autocorrelation (M. W. Woolrich et al., 2001) correction to assess task-evoked changes in neural responses to social and reward stimuli employing two main models. Both models employed a general linear model (GLM) approach, convolved using the canonical hemodynamic response function. Each first level GLM included two regressors to account for the block structure in a task: in the gambling task, these regressors corresponded to reward and loss, and in the social task, mental trials and random trials. The reward regressor was modeled from the onset of the first trial of the block, which consisted of mostly reward until the offset of that block; similarly, the punishment regressor was elicited the same way, but drawn from the mostly-loss blocks. The mental regressor was modeled from the onset of the first block of the run that consisted of mostly mental videos until the offset of that run; the random regressor was similarly derived from the run that consisted of mostly random trials.

To investigate task-evoked changes in brain connectivity, we utilized a psychophysiological interaction (PPI) analysis (Friston et al., 1997). Our PPI models focused on cerebellar coupling with vmPFC and were later expanded in an exploratory whole brain analysis, using the vmPFC as the seed region. The vmPFC region of interest (ROI) was defined based on a preliminary analysis using FLAME. For this analysis, we thresholded and corrected for multiple comparisons within a priori ROI (http://aspredicted.org/blind.php?x=8qw2h3) at the voxel level. ROIs were identified via a Cerebellar Atlas in MNI152 space after normalization with FNIRT and the Harvard-Oxford cortical and subcortical atlases in FSL. In addition to the same regressors used in the activation models, our PPI models included regressors to represent the activation time course in connectivity between the vmPFC seed region and cerebellar target regions. Similar to our analysis of activation, we evaluated two distinct PPI models for each task: the social cognition and reward processing tasks.

To combine data across runs for our second-level analysis, we used a fixed effects model, with random effects variance forced to zero in FLAME (Beckmann et al., 2003; M. Woolrich, 2008; M. W. Woolrich et al., 2004). Next, we combined data across participants using FLAME (FMRIB’s Local Analysis of Mixed Effects) stage 1 (Beckmann et al., 2003; M. Woolrich, 2008; M. W. Woolrich et al., 2004). To detect group differences in neural activity for the social cognition and reward processing tasks, and to control for other factors that influence our results, we used a GLM consisting of 11 regressors. The 11 regressors contained data describing participant head movement during the task, gender, depressive symptoms, sadness scores, impulsivity, alcohol and tobacco use. Gaussianized z-statistic images were thresholded with voxel-level GRF-theory-based maximum height thresholding with a corrected significance threshold of *p*=0.05 (Worsley, 2001). Our model also included F-tests to directly evaluate our ability to reject the null hypothesis that MDD = FH = HC with regard to each of our hypotheses. We controlled for family structure in the data (Winkler et al., 2015) using PALM (Winkler et al., 2014) to evaluate pre-registered hypotheses regarding group differences in behavior and neural responses tied to activation and connectivity. We note, however, that the seed for our connectivity analysis (vmPFC) was identified through conventional parametric statistics since the goal of that analysis was simply to identify regions that respond similarly to social and reward, irrespective of group. Our application of PALM was one-tailed, and we controlled for the number of comparisons across contrasts using the -corrcon flag, which includes FWER-correction of p-values. We also enabled threshold-free cluster enhancement (TFCE) inference for univariate tests, and was permuted 5000 times for all behavioral, activation, and PPI analyses.

In addition, we performed t-tests between MDD, FH, and HCs to determine whether the groups differed by head motion variables in both the gambling and social cognition task. We found no difference among groups on the gambling task between MDD and FH (*t*(172)=-1.28, *p*=0.20), MDD and HC (*t*(174)=-0.13, *p*=0.88), FH and HC (*t*(206)=1.26, *p*=0.20), nor any difference among groups on the social cognition task between MDD and FH (*t*(172)=-0.99, *p*=0.31), MDD and HC (*t*(174)=0.01, *p*=0.99), FH and HC (*t*(206)=1.15, *p*=0.24).

## Results

### Behavioral and Clinical Analyses

Our analyses began by testing whether a FH of depression was associated with higher depression symptoms and sadness experienced across one’s lifetime. To compare depression symptoms, we evaluated the average number of lifetime depressive symptoms reported by each participant who met DSM-IV criteria. We found that individuals with a FH reported more lifetime depressive symptoms across their lives compared to HCs (*t*(206)=2.54, *p*=0.011). Individuals in the depression group had higher depressive symptom rates than both FH (*t*(172)= 19.22, *p*<0.01) and HC (*t*(174)=27.00, *p*<0.01; see Figure 1A) groups. Similarly, we used unadjusted sadness scores as measured by the NIH Toolbox Sadness Survey to compare self-reports of sadness across groups. The NIH Toolbox Sadness Survey is a subcomponent of a larger battery, the NIH Toolbox Emotion Battery, a standardized form of measuring emotional health across the lifespan (www.nihtoolbox.org). We found that unlike the depressive symptoms, sadness scores in participants with a FH of depression did not differ significantly from HCs (*t*(206)=1.88, *p*=0.06). However, as expected, the levels of sadness reported by the depression group surpassed both FH (*t*(172)=4.39, *p*<0.01) and HC groups (*t*(174)=6.09, *p*<0.01, See Figure 1b). Additionally, we tested for differences in alcohol and tobacco use (in the past 7 days) between groups, as previous work has found a relationship between tobacco use (Edwards et al., 2011) or occasional bouts of heavy drinking (Manninen et al., 2006) and depression. However, we did not find significant differences in either alcohol or tobacco use across groups (see Table 1).

**Table 1:** Percentage of male and female participants, as well as average tobacco (measured in number of cigarettes) and alcohol (measured in number of drinks) used in the past 7 days across each group. Groups did not differ significantly in their use of alcohol over the past 7 days. Alcohol use in MDD and FH (*t*(172)=0.47, *p*=0.63, *d*=0.07), MDD and HCs (*t*(174)=0.34, *p*=0.73, *d*=0.05), FH and HCs (*t*(206)=-0.15, *p*=0.88, *d*=-0.02). Similarly, there were no differences in tobacco use between MDD and FH (*t*(172)= −0.36, *p*=0.71, *d*=-0.05), MDD and HCs (*t*(174)= 0.12, *p*=0.90, *d*=0.01), and FH and HCs (*t*(206)=0.51, *p*=0.61, *d*=0.07).

| Group | Participants (N) | Gender | Alcohol Use (drinks/7 days) | Tobacco Use (cigarettes/7 days) |
| --- | --- | --- | --- | --- |
| Depression (MDD) | 71 | 61% F; 40% M | 4.661 | 5.450 |
| Family history (FH) | 103 | 62% F; 38% M | 4.165 | 6.533 |
| Healthy controls (HC) | 105 | 50% F; 50% M | 4.295 | 5.123 |

**Figure 1:**
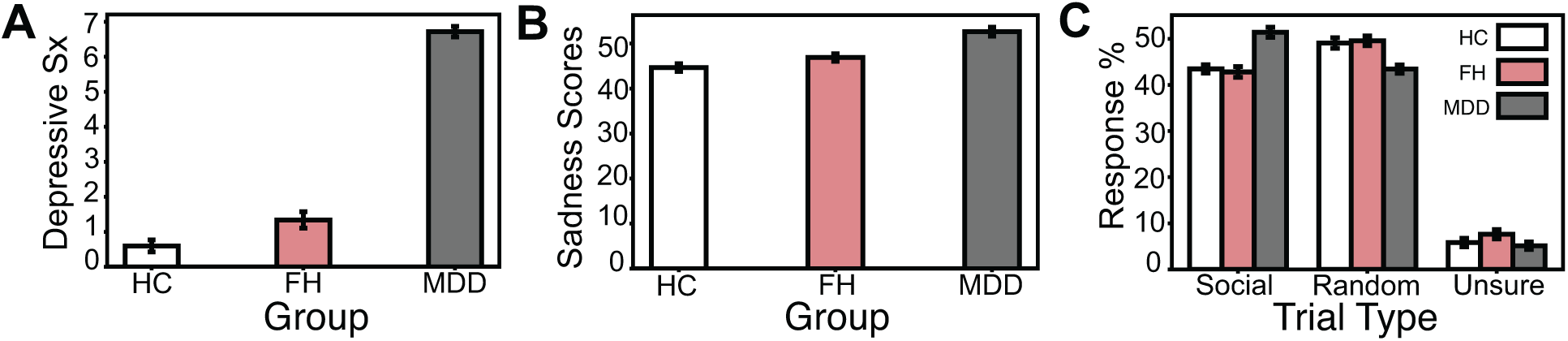
Self-reported depression symptoms and levels of sadness across depression, family history, and healthy control groups. ***A***, Depression (MDD) group shows higher average depression symptoms compared to both individuals with a family history of depression (FH; *t*(172)= 19.22, *p*<0.01, *d*=2.82) and healthy controls (HCs; *t*(174)=27.00, *p*<0.01. *d*=24.00). Individuals in the FH group show higher depression symptoms compared to HCs (*t*(206)=2.54, *p*=0.011, *d*=0.35). ***B***, Unadjusted sadness scores are significantly higher in MDD when comparing to both FH (*t*(172)=4.39, *p*<0.01, *d*=0.68) and HC groups (*t*(174)=6.11, *p*<0.01, *d*=0.93), but did not reach our significance threshold between FH and HCs (*t*(206)=1.88, *p*=0.06, *d*=0.26). ***C***, Average social, random, and unsure percent responses to a Theory of Mind (ToM) social cognition task presented to participants during the fMRI scanning paradigm. Initially, MDD showed a higher percentage of ‘social’ selections relative to FH (*t*(172)=5.81 *p*<0.01, *d*=0.86) and HC (*t*(174)=5.66, *p*<0.01, *d*=0.83) groups. MDD also initially demonstrated lower percentages of ‘random’ selections relative to FH (*t*(172)=-3.89 *p*<0.01, *d*=-0.59) and HCs (*t*(174)=-3.87, *p*<0.01, *d*=-0.59). MDD and FH (*t*(172)=-1.81 *p*=0.07), MDD and HCs (*t*(174)=-0.57, *p*=0.57, *d*=-0.08), FH and HCs (*t*(206)=1.41, *p*=0.16, *d=*0.19) did not differ significantly with respect to percentage of ‘unsure’ selections throughout the task. However, after controlling for family structure in the data, any differences in performance between groups on the social cognition task were eliminated (all −log10(*p*) values < 1.3010, where *p*=0.05, corrected for multiple comparisons across voxels and contrasts).

We hypothesized that the depression group would be more unsure about the social nature of the stimuli relative to FH or HC groups. To test this hypothesis, we compared the selections each group made while engaging in the ToM task in the scanner. Contrary to our expectations, we initially found that the depression group had a higher percentage of ‘social’ selections relative to FH (*t*(172)=5.81 *p*<0.01) and HC (*t*(174)=5.66, p<0.01) groups. The depression group also initially demonstrated lower percentages of ‘random’ selections relative to FH (*t*(172)=-3.89 *p*<0.01) and HC (*t*(174)=-3.87, *p*<0.01) groups. Depression and FH (*t*(172)=-1.81 *p*=0.07), depression and HCs (*t*(174)=-0.57, *p*=0.57), FH and HCs (*t*(206)=1.41, *p*=0.16) did not differ significantly with respect to percentage of ‘unsure’ selections throughout the task (see Figure 1c). However, after controlling for family structure in the data, we found no differences in performance between groups on the social cognition task (all −log_10_(*p*) values < 1.3010, where *p* is 0.05, corrected for multiple comparisons across voxels and contrasts).

### ROI Analyses

Next, we evaluated whether the social cognition and reward processing tasks activate canonical mentalizing and reward processing regions respectively, and if activity overlaps. We found that independent of group, the social cognition task activated the TPJ on the mental > random trial contrast, and the reward processing task activated the nucleus accumbens on the reward > punishment trial contrast. A subsequent conjunction analysis revealed that a region of the vmPFC responded to both tasks, independent of groups (Figure 2). We then used this region of the vmPFC as a seed for psychophysiological interaction (PPI) analyses to evaluate vmPFC connectivity patterns with the cerebellum and later, across the entire brain.

**Figure 2:**
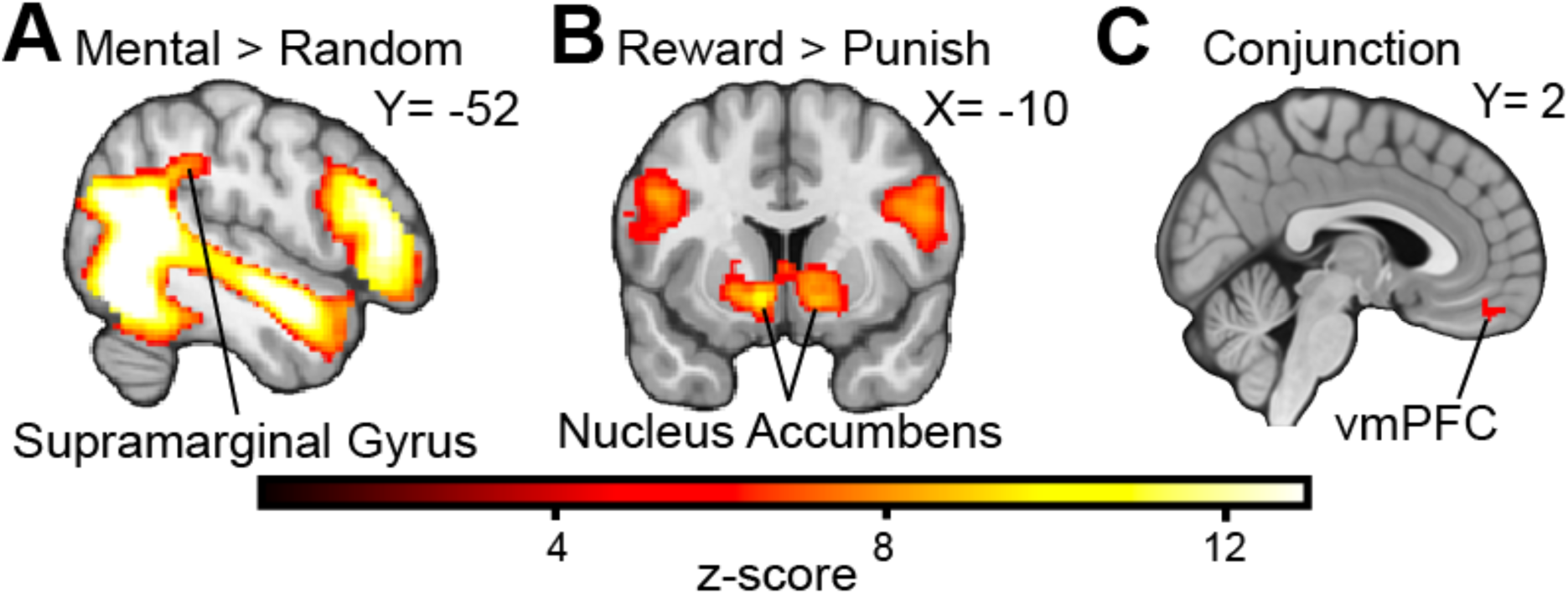
Activation and Conjunction Analyses. ***A***, Independent of group, the social cognition task evoked activity in the temporoparietal junction specific to Mental > Random trials contrasts. ***B***, Similarly, the reward processing task evoked striatal activation on the Reward > Punishment trial contrasts. ***C***, Both tasks also demonstrate shared activation in a region of the ventromedial prefrontal cortex (vmPFC). Gaussianized z-statistic images were thresholded with voxel-level GRF-theory-based maximum height thresholding with a corrected significance threshold of p=0.05.

We then evaluated the presence of dose-dependent responsivity among the groups during the social cognition task. We predicted that during this task, MDD would demonstrate blunted TPJ activation, but increased cerebellar activity, particularly in the vermis and posterior hemispheres relative to HC and FH groups. Our dose-dependency hypotheses also anticipated that while the MDD group would show the largest responses (either by way of increase or decrease), the FH group would trend towards the pattern exhibited by the depression group, with blunted TPJ activation and increased cerebellar activity relative to HCs. Additionally, we expected to see that the MDD group would have blunted striatal activation in response to the reward processing task, relative to HC and FH groups, while the FH group also would show blunted striatal activation relative to HCs. Despite these hypotheses, we found no differences in TPJ or cerebellar activation within the social task, nor striatal activation differences during the reward processing task across the groups.

We also expected to see decreased vmPFC-cerebellar (specifically with respect to the bilateral lobule VIIb) connectivity during the social cognition task for the MDD group relative to FH and HC groups. As with our activation hypotheses, we expected that the FH group also would show decreased connectivity between the vmPFC and cerebellar subregions, albeit relative to HCs. Our results do not show any differences in vmPFC-cerebellar functional connectivity between MDD and FH groups, but are seen when comparing MDD and HC groups, such that MDD groups show a small region of increased vmPFC-cerebellar connectivity during the mental condition (see Table 2). Nevertheless, we find increased vmPFC-cerebellar functional connectivity in the FH group relative to HCs. Specifically, relative to HCs, the FH group shows increases in connectivity between the vmPFC and the left crus I of the cerebellum during the mentalizing trials. Random trials, on the other hand, also show increased connectivity between the vmPFC and left crus I, as well as the left VIIb region of the cerebellum, with the FH group again showing greater connectivity than the HCs in particular (see Figure 3).

**Table 2:** vmPFC Seed Functional Connectivity by task, condition, and group comparison. Regions that demonstrate significant activation above the −log10(*p*) value cutoff of 1.3010 (where *p* is 0.05, corrected for multiple comparisons across voxels and contrasts) among Reward (Rew), Punishment (Pun), Mental, and Random conditions for the Gambling and Social tasks, respectively. Sites of functional connectivity activation marked with probabilistic anatomical labels correspond to anatomical coordinates (x,y,z) for each group comparison: major depressive disorder (MDD), family history (FH) and healthy controls (HC).

| **Table 2: vmPFC Seed Functional Connectivity by task, condition, and group comparison** |  |  |  |  |  |  |  |
| --- | --- | --- | --- | --- | --- | --- | --- |
| **vmPFC-Cerebellar PPI analyses** |  |  |  |  |  |  |  |
| **Task** | **Condition** | **Comparison** | **Probabilistic Anatomical label** | **x** | **y** | **z** | **-log<sub>10</sub>(p)** |
| **Gambling** | Rew>Pun | MDD>FH | Right Crus I (15%), Right Crus II (5%) | 9 | -84 | -20 | 1.6307 |
| **Gambling** | Pun>Rew | FH>MDD | Right Crus I (15%), Right Crus II (5%) | 9 | -84 | -20 | 1.6307 |
| **Social** | Mental | MDD>HC | Right VIIa (45%), Right VIIb (45%) | 14 | -72 | -54 | 1.6497 |
| **Social** | Mental | FH>HC | Left Crus I (100%) | -47 | -68 | -36 | 1.6055 |
| **Social** | Random | FH>HC | Left Crus I (71%), Left VI (28%) | -36 | -61 | -28 | 1.6126 |
| **vmPFC-whole brain PPI analyses** |  |  |  |  |  |  |  |
| **Social** | Mental | FH>HC | Insular Cortex (39%) | -30 | 13 | 6 | 1.4294 |
|  |  |  | Left Putamen (98%) | -17 | 9 | -7 |  |
|  |  |  | Frontal Pole (78%) | -36 | 42 | 25 |  |
| **Social** | Random | FH>MDD | Frontal Pole (90%) | -42 | 53 | -3 | 1.3851 |

**Figure 3:**
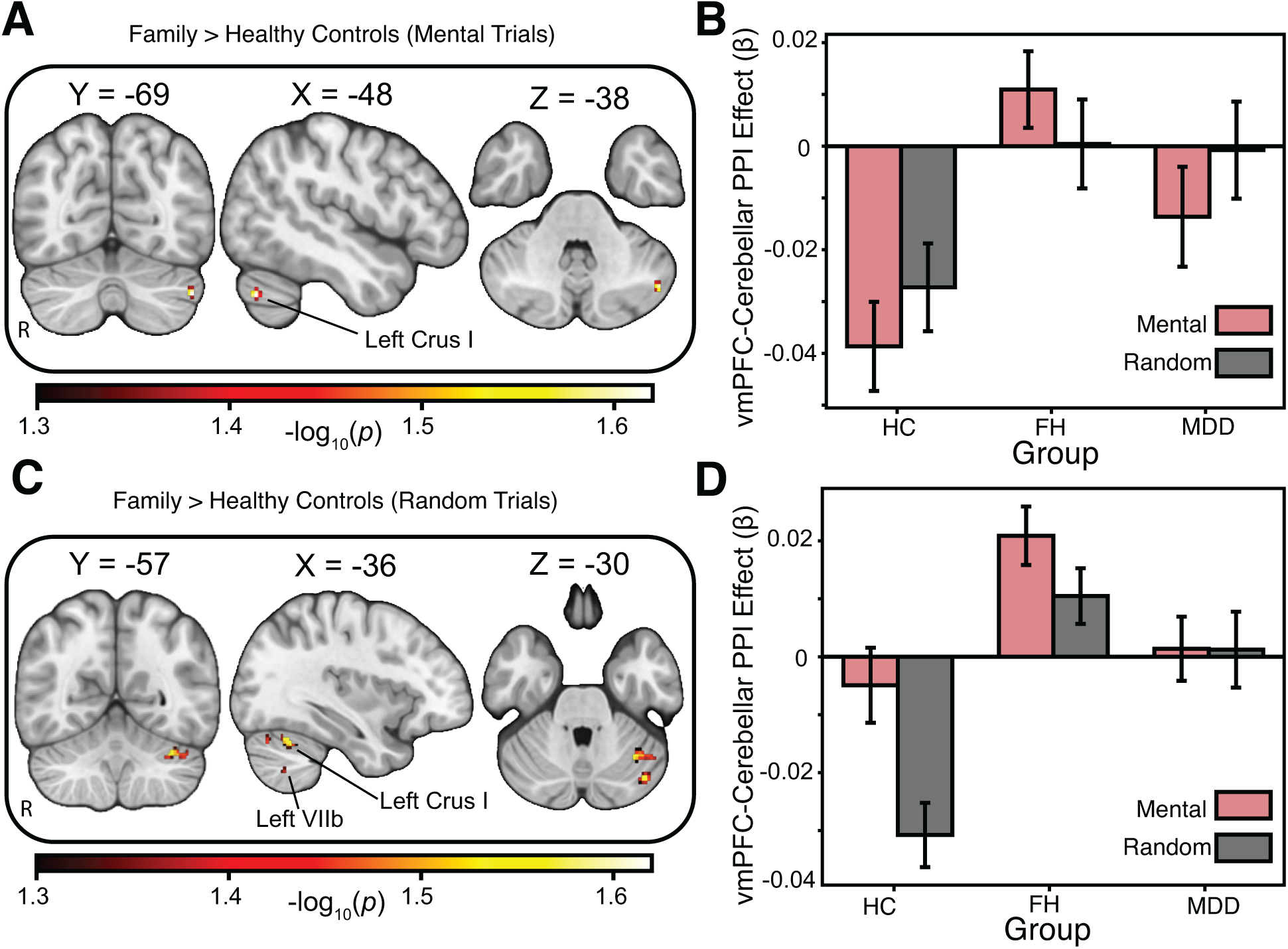
Psychophysiological (PPI) analysis using vmPFC as the seed region. ***A***, Relative to healthy controls, the family history group showed increases in task-based functional connectivity between the vmPFC and the left crus I of the cerebellum during the “Mentalizing” trials. Areas of activation passed an a priori small volume correction (cerebellum) using threshold-free-cluster enhancement (TFCE) and permutation-based testing using PALM. ***B***, Interrogation of the left crus I region of the cerebellum revealed increased connectivity relative to the decrease in activation seen in healthy controls during the mental trials. ***C***, Increased connectivity between the vmPFC and left crus I and left VIIb regions of the cerebellum was selective for the random stimuli in the social task, with the family history group showing a greater response than the healthy controls. Areas of activation passed an a priori small volume correction (cerebellum) using threshold-free-cluster enhancement (TFCE) and permutation-based testing using PALM. ***D***, Interrogation of the left crus I and left VIIb regions of the cerebellum showed the family history group had a larger increase in connectivity relative to healthy controls during the mental trials.

### Exploratory Analyses

We expanded our search beyond the pre-registered hypotheses and a priori cerebellar target to include a whole-brain analysis of vmPFC connectivity. We again find functional connectivity differences primarily for the FH and HC groups during the social cognition task. Specifically, individuals with a familial risk of depression, relative to healthy controls, demonstrated increased vmPFC-nucleus accumbens and vmPFC-dorsolateral PFC functional connectivity during the mentalizing trials (see Figure 4).

**Figure 4:**
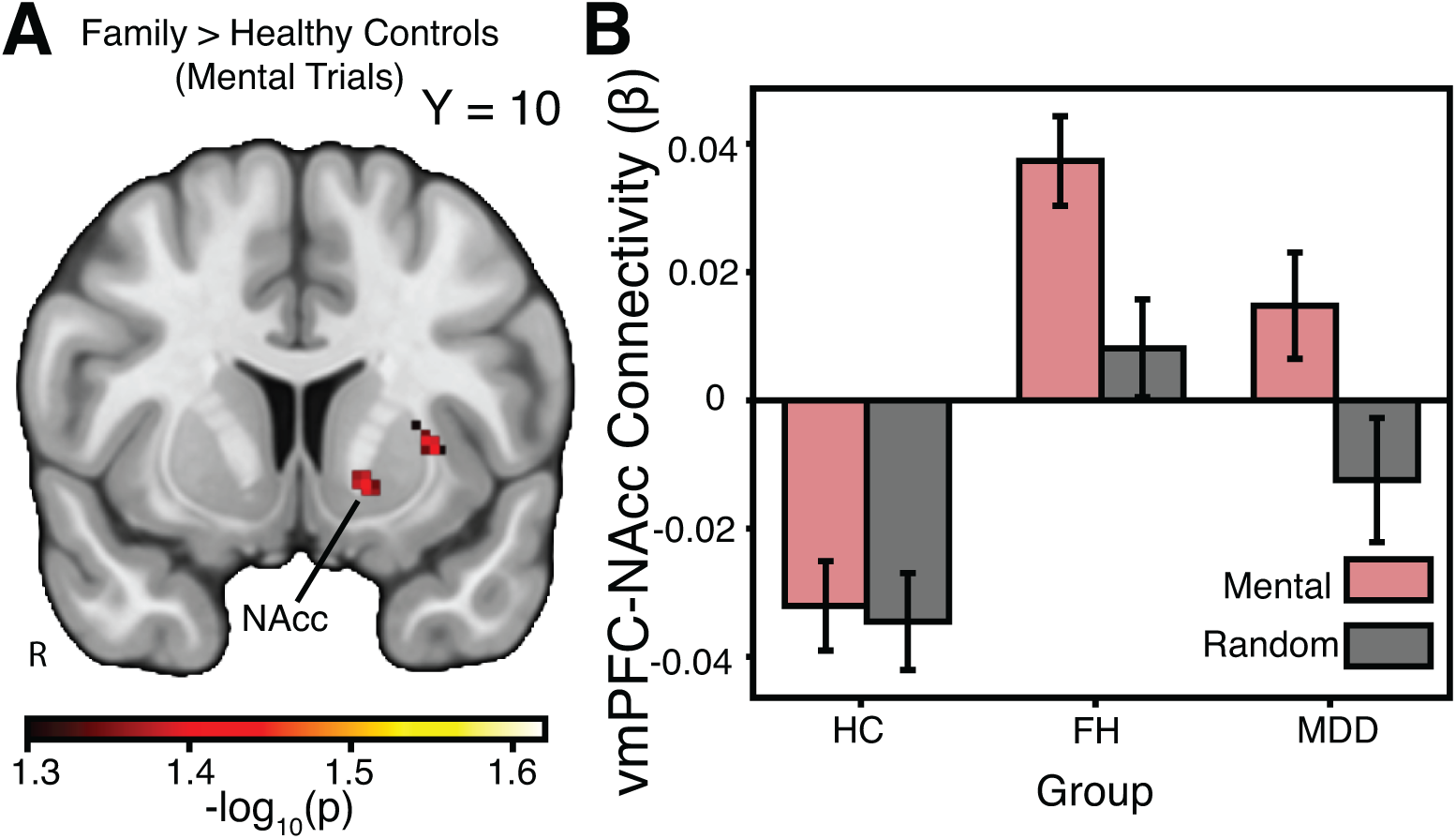
Exploratory whole-brain analysis. A, Relative to healthy controls, the family history group showed greater task-based functional connectivity between the vmPFC and the nucleus accumbens (NAcc) during the mental trials of the social cognition task. Areas of activation passed a whole brain correction using threshold-free-cluster enhancement (TFCE) and permutation-based testing. B, Interrogation of the NAcc revealed increased connectivity in the family history group, whereas the healthy control group demonstrated a relative decrease in connectivity during the mental trials.

## Discussion

A deeper understanding of the neural mechanisms underlying social cognition and reward processing can elucidate how these systems typically function, and the conditions under which they are susceptible to change. Mapping task-based neural activity among those with a well-established risk factor for MDD allows us to probe this susceptibility to change in reward and social systems. Moreover, capturing the state of the brain before MDD onset in an at-risk group is especially relevant to clinical efforts in treatment and intervention. Thus, the goal of the current study centered on evaluating task-based neural processing of social and reward systems in those with a FH of major depressive disorder, using data from the Human Connectome Project. In line with our expectations, we found that relative to HCs, individuals with a FH of MDD demonstrate increases in depressive symptoms, and the MDD group showed increased sadness scores and depressive symptoms relative to both FH and HC groups. Yet, we did not find significant neural differences between HCs and those with a personal history of MDD on either social or reward processing tasks as expected. Nevertheless, we confirmed our initial hypothesis that the FH group would show neural alterations relative to HCs during the social cognition task. Specifically, we found that relative to HCs, individuals with a FH of MDD demonstrated increased task-dependent functional connectivity between the vmPFC and the nucleus accumbens, left dorsolateral PFC, and subregions of the cerebellum during a mentalizing task. Altogether, these results reveal that in addition to upticks in pre-clinical reports of depressive symptoms, individuals with a FH of depression may be at risk for neural abnormalities underlying ToM processes, with altered cerebellar-vmPFC functional connectivity contributing to this difference.

A host of recent studies have begun to reveal the functional role of the cerebellum in social, cognitive, and affective neuroscience (Van Overwalle et al., 2014, 2015). For example, the cerebellum has been linked to social reward (Carta et al., 2019), social preference (Badura et al., 2018), and emotional processing (Baumann & Mattingley, 2012). Of particular relevance to our current results, the cerebellum has been linked to mentalizing processes by way of connectivity with the cortex, including the TPJ (Van Overwalle et al., 2019) and dorsomedial PFC (Van Overwalle et al., 2015). Cerebellar connectivity also has been tied to the default mode network (Buckner et al., 2011), which participates in social inference (Buckner & DiNicola, 2019). In light of this past work, the present study dovetails with the existing literature by providing additional evidence demonstrating involvement of cerebellar connectivity during mentalizing. Specifically, our findings implicating task-based vmPFC-cerebellar connectivity differences in a ToM task appears to fall in line with previous suggestions that the cerebellum is involved in social reasoning (Van Overwalle et al., 2019).

Unlike prior studies of the cerebellum, our work highlights a novel role for the region’s involvement in the vulnerabilities inherited through a FH of depression. Whereas investigations of autism (Becker & Stoodley, 2013), schizophrenia (Andreasen & Pierson, 2008), and major depressive disorder (Liu et al., 2012) have shown alterations in the cerebellum related to social and affective processing, these data demarcate populations that already have developed a disorder. As such, relatively little is known about the neural signatures involved in a FH of depression as it relates to social cognition. Through the identification of altered task-based cerebellar functional connectivity with the vmPFC, we shed new light on possible mechanisms at play that may lead to the early stages of MDD. In addition to building a deeper understanding of the cerebellum’s role in social cognition, this work may inform interventions geared towards mitigating later emergence of MDD. Indeed, considering the cerebellum’s influence on social and cognitive development, the region previously has been suggested as a potential site for treatment (S. S. H. Wang et al., 2014). Moreover, altered functional connectivity between the cortex and cerebellum reliably can predict the presence of depression (Ma et al., 2013). The results of the present study indicate that consideration of the cerebellum as a therapeutic target may be promising early on, before the onset of MDD.

Although our study demonstrates that task-dependent connectivity with the cerebellum is altered in those with a FH of depression, we note that our findings are accompanied by some limitations. First, we note that the current work uses a block design and our analyses focus on differences between blocks, consistent with other work using the HCP data (e.g., Barch et al., 2013). Yet, some studies using the HCP data have suggested that the reward task also could be analyzed based on individual events, which may help separate processes tied to valence and salience (Zhang et al., 2017). Future studies may be able to build on our findings by examining salience and valence and by considering other datasets that are optimized for such questions (e.g., ABCD). Additionally, our results that vmPFC-cerebellar connectivity was enhanced in the FH group relative to HCs was not specific to the mental or random conditions in the social task. We speculate that the lack of differences between the two conditions in the social task could be due to the nature of the task. Indeed, although the task used by the HCP is a robust probe of TPJ activation, it may not evoke similar levels of activation in other regions that process more subtle differences in how participants process and interpret a range of social interactions. Future work building on these findings would do well to involve measures of social cognition that characterize mental representations of others in a variety of ways. For instance, it may be particularly relevant to examine social interactions as seen in the virtual school paradigm (Jarcho et al., 2016), as this may better probe how the cerebellum processes the mental states of characters in a more ecologically valid setting.

In addition to limitations with the social task, we also note that our study may not be able to speak directly to the mechanisms underlying risk for developing MDD, given the cross-sectional nature of the Human Connectome Project. Although our original hypothesis—that aberrant patterns of connectivity would deviate further from HCs when examining people with a personal history of depression—was not supported by our findings, we note that this is an area where longitudinal data would be necessary for understanding whether and how altered connectivity is associated with the development of MDD. To address this question, future research could leverage other open datasets that include longitudinal assessments of cerebellar connectivity, such as the Adolescent Brain Cognitive Development study (ABCD; (Casey et al., 2018). Thus, in the absence of longitudinal data, it is unclear whether neural differences between FH and HC groups can predict a subsequent MDD diagnosis.

Despite these limitations, our study demonstrates that relative to HCs, individuals with a FH of MDD exhibit altered task-dependent vmPFC-cerebellar activity during a social cognition task. Previous research shows that the cerebellum plays a key role in processing social, affective and rewarding stimuli and is a region of interest in psychiatric illnesses, including MDD (Konarski et al., 2005). Evidence for neural alterations underlying social cognition in advance of a personal episode of MDD implies the possibility of early intervention and support that may mitigate later emergence of the disorder. Abnormal neural connectivity has been documented in subclinical depression in another study using the HCP dataset (Ely et al., 2016), as well as other disorders (Young et al., 2015) and may play a central role in the development of psychiatric illnesses (Zhang et al., 2015) and maladaptive behaviors (Diehl et al., 2018) such as substance use (Cheng et al., 2019). From a clinical lens, the identification of neural alterations and their ability to establish a diagnosis or predict how an individual will respond to treatment remains unclear (Savitz et al., 2013). Yet, as research accumulates data detailing the mechanisms driving neural alterations, future work may more reasonably employ emergent technology—such as brain stimulation—to intervene (Diehl et al., 2018). Considering the limits MDD places on social cognition (Knight et al., 2018), addressing abnormalities in the underlying neural mechanisms of social cognition in particular lends itself to the development of treatments that facilitate social support and connection, a leading protective factor against future depressive episodes (Cruwys et al., 2014). Additional work must continue to investigate how the cerebellum connects with other regions across a variety of affective disorders to understand the full spectrum of the site’s role in healthy and abnormal social cognition alike.

## Data Availability Statement

Data were provided [in part] by the Human Connectome Project, WU-Minn Consortium (Principal Investigators: David Van Essen and Kamil Ugurbil; 1U54MH091657) funded by the 16 NIH Institutes and Centers that support the NIH Blueprint for Neuroscience Research; and by the McDonnell Center for Systems Neuroscience at Washington University. This study was pre-registered, and can be found at http://aspredicted.org/blind.php?x=8qw2h3. Study materials and code is available on Open Science Framework (DOI 10.17605/OSF.IO/JU32V). Images are available on the NeuroVault repository at https://identifiers.org/neurovault.collection:6130.

## Acknowledgments

This work was supported, in part, by grants from the National Institutes of Health (R21-MH113917 and R03-DA046733 to DVS) and the American Association for University Women (Career Development Grant to LJT). DVS was a Research Fellow of the Public Policy Lab at Temple University during the preparation of this manuscript (2019-2020 academic year). We thank Ka-Yi Chat for helpful feedback on the manuscript.

